# Resequencing the *Escherichia coli* genome by GenoCare single molecule sequencing platform

**DOI:** 10.1101/163089

**Authors:** Luyang Zhao, Liwei Deng, Gailing Li, Huan Jin, Jinsen Cai, Huan Shang, Yan Li, Andrew X. Yang, Fang Chen, Zhi Zhao, Guanjie Xu, Wuxing Liu, Siyu Liu, Guobing Xiang, Bin Liu, Weibin Xu, Lidong Zeng, Renli Zhang, Huan Zhao, Ping Wu, Zhiliang Zhou, Jiao Zheng, Pierre Ezanno, Weiyue Chen, Qin Yan, Michael W. Deem, Jun Yu, Jiankui He

## Abstract

Next generation sequencing (NGS) has revolutionized life sciences research. Recently, a new class of third-generation sequencing platforms has arrived to meet increasing demands in the clinic, capable of directly measuring DNA and RNA sequences at the single-molecule level without amplification. Here, we use the new GenoCare single molecule sequencing platform from Direct Genomics to resequence the *E. coli* genome and show comparable performance to the Illumina MiSeq system. Our platform detects single-molecule fluorescence by total internal reflection microscopy, with sequencing-by-synthesis chemistry. With a consensus sequence of 99.71% nucleotide identity to that of the Illumina MiSeq system’s, GenoCare was determined to be a reliable platform for single-molecule sequencing, with strong potential for clinical applications.

## INTRODUCTION

Since the primer-extension strategy developed by Frederick Sanger in the 1970s, based on chain-terminating modified nucleotides [1], next-generation sequencing (NGS) technologies have revolutionized life sciences research. The first two publications of the human genome sequence [2][3] are still considered major milestones today. DNA sequencing has revolutionized biological investigations in basic science as well as clinical diagnosis [4] [5]. Some applications in the emerging field of precision medicine [6] include cancer diagnosis [7] [8] and inherited disease diagnosis [9] [10]. Progress on NGS technologies in medicine also introduces many ethical questions [11] [12]. Besides human health care, other promising applications of sequencing technologies include detection of pathogenic organisms [13] [14] and DNA profiling in forensic sciences [15] [16].

Regarding genomics, NGS technologies serve two goals: genome mapping and de-novo assembly. These two goals mainly differ in the data processing downstream of the sequencing step [17]. Genome mapping aims to determine the sequence of a given genome by comparison with an existing consensus sequence [18], and can be achieved by performing massively parallel sequencing of short reads (i.e. 25-40 bases). De-novo assembly consists of sequencing a novel genome without a consensus sequence, and generally requires longer reads and more computing capacities than does the mapping approach [19] [20].

A focus on genome mapping, simple operation, cost-effective sample preparation, high throughput data generation, and better instrument sensitivity is key to the human genome sequencing market in the near future. Therefore, improving instruments is an important necessity. Currently, the cost of the sample preparation for NGS is still a significant part of the total cost of genome sequencing. Moreover, the final cost for sequencing human genomes in a wide group of individuals is still hardly affordable no matter the technology used. Single molecule sequencing was first experimented with in the late 1980s [21]. This technological breakthrough is now seen as the next step beyond NGS, contributing to the advent of very sensitive instruments [22] [23].

Different SM sequencing technologies have rapidly developed over the past decade, with progress on read length, sequencing time, and data throughput. Three technologies are now well known, each with their unique characteristics: (i) the first true single molecule sequencing (tSMS) combined with sequencing-by-synthesis (SBS) [24] technology from Helicos Biosciences [25, 26]; (ii) single molecule real time (SMRT) sequencing technology from Pacific Biosciences producing super long read length (longer than 10k bases [27, 28]), but relatively low throughput; and (iii) Oxford Nanopore Technologies, producing long read length (6k bases [27]) but limited accuracy and low throughput. The GenoCare platform improves on principles from the Helicos Biosciences platform.

A combination of minimal, amplication-free sample preparation and efficient massively-parallel short reads processing are ideal for the demands of sequencing-based clinical diagnosis. Advantages of GenoCare single molecule sequencing include (i) a simple and time-saving sample preparation consisting of DNA shearing followed by poly-A tailing and 3' end blocking steps, (ii) absence of PCR amplification, and (iii) potential for direct RNA sequencing for investigation of transcriptomic aspects of gene expression.

Our approach is devised to provide simple operation and high-throughput, unbiased data. Recently, we demonstrated a direct targeted sequencing of cancer related gene mutations [37] and M13 virus genome sequencing [38] at the single molecule level. In this study, we resequenced the *E. coli* genome using two different platforms: the Illumina MiSeq system and the new Direct Genomics GenoCare platform. We describe the potential of GenoCare as a single molecule sequencing platform for the clinic.

## MATERIALS AND METHODS

### Sample Preparation

#### E. coli Sample

The *E. coli* sample was purchased from Affymetrix as product #14380. The strain is ATCC 11303.

#### Oligonucleotide Primers

5’ amine functionalized Poly-T oligonucleotides were purchased from Sangon and used as received.

#### DNA Fragmentation

*E. coli* DNA was randomly fragmented into ~200bp dsDNA fragments using NEBNext^®^ dsDNA Fragmentase^®^ (from NEB, ref M0348S). These DNA fragments were then purified using Agencourt AMPure XP beads (from Beckman, ref. A63881). DNA concentration was assessed by UV absorption using a Nanodrop 2000 device.

#### Poly-A Tailing and Blocking

Multiple incorporations of 50-100 dATP at the 3' end of ssDNA fragments resulted in a poly-A tail; this reaction completed within 20 minutes. Poly-A tailed 3' ends were then blocked by incorporation of Cyanine 3 dideoxy ATP (Cy3-ddATP from PERKINELMER, ref. NEL586001EA). This blocking reaction completed within 30 minutes using the enzyme Terminal Transferase (from NEB, ref. M0315), preventing incorporation of reversible terminators at the 3' end of the template strands.

### Surfaces and Template Capture

#### Surface Chemistry

Sequencing surfaces were prepared on 110×74 mm epoxy-coated glass coverslips (SCHOTT, Jena, Germany). Poly-T oligonucleotides were covalently bond to surface.

##### Flow Cells

The functionalized glass coverslip was assembled with a 1.0 mm thick glass slide by a pressure sensitive adhesive to form a flow cell. The flow cell has 16 channels, determined by the adhesive shape. For *E. coli* sequencing in this experiment, ~0.5% of one channel was imaged.

##### Template Capture (Hybridization)

The surface of the flow cell was chemically modified by anchoring poly-T ssDNA strands at their 5' end. This anchoring allows for capture of poly-A tailed strands from the library once injected inside the flow-cell at 55 °C. Non-hybridized templates were then washed away by 150 mM HEPES, 1X SSC and 0.1% SDS, followed by 150 mM HEPES and 150 mM NaCl.

#### Sequencing Reactions

##### The GenoCare Platform

Our sequencing tests have all been performed on the GenoCare platform designed and fabricated by the Direct Genomics company. The GenoCare sequencer has been developed on the basis of a single molecule sequencing-by-synthesis process described previously [37][38]. The platform is highly automated and user-friendly. It comprises three major parts: the fluorescence imaging system, the microfluidic system, and the automated sample stage. Fill & Lock was done before sequencing started. To acquire the sequencing results reported here, 120 sequencing cycles were carried out, and the total experimental time was *ca.* 23 hours including sample preparation time (*ca.* 3 hours) and GenoCare sequencing time (*ca.* 20 hours). During each sequencing cycle, only one type of terminator was added to the flow-cell for nucleotide incorporation. Each quads (4 cycles) of terminator addition follows the order of CTAG. The flow-cell channels were then treated with a rinse procedure so that the free terminators not incorporated in any strand could be flushed out. The flow-cell was subsequently fluorescence-imaged using total internal reflection fluorescence (TIRF) microscopy, followed by fluorophore cleavage and residual bond capping procedures. The integration time for each field of view (FOV) during the imaging process has been optimized to 200 ms, which provides satisfactory signal-to-noise ratio while keeping the photobleaching of dyes at a negligible level.

#### Alignment

##### Read Filtering

Reads with more than 8 consecutive cycle repeats (CTAG) were filtered out before alignment.

##### Read Alignment

Bowtie 2 software [39] was used for all read alignment. The following options were applied: -local -D 25 -R 3 -N 1 -L 18 -i S,1,0.50. The K-12 genome was used as reference. Because the true reference was unavailable, to compare Illumina and GenoCare similarity, we aligned GenoCare reads to the consensus sequence of Illumina reads aligned to the K-12 genome as reference.

##### Analysis

Python scripts, Samtools [40], GATK [41], and Qualimap 2 [42] were used to generate alignment statistics. Average identity of consensus sequences compared with reference sequences was calculated with the dnadiff tool from MUMmer [43]. Graphs were produced with the R package ggplot2 [44].

### RESULTS AND DISCUSSION

#### Alignment Statistics

Because the reference genome for the *E. coli* sample used was not available,we used the common K-12 genome as reference. For the GenoCare platform, 2,329,974 reads (31.33%) were uniquely aligned to the K-12 reference genome, with 91.39% of genome covered (**Table 1**). Using the MUMmer package’s dnadiff tool, pairwise comparisons of the alignment’s consensus sequence with the K-12 reference sequence found 98.97% average nucleotide identity. For comparison, the Illumina platform yielded 90.60% of genome covered and 98.73% average nucleotide identity to the K-12 genome. In order to demonstrate reliability of GenoCare reads in spite of lacking the original reference sequence, we aligned GenoCare reads to the consensus sequence obtained by aligning Illumina reads to the K-12 genome as a reference. This alignment gave average identity of 99.71%, indicating comparable and reliable sequencing data generated by the GenoCare platform.

**Table 1.** Alignment Statistics for uniquely mapped reads.

|  | Illumina<br>(K-12 ref.) | GenoCare<br>(K-12 ref.) | GenoCare<br>(Illumina consensus ref.) |
| --- | --- | --- | --- |
| **Mean Coverage** | 84.71X | 14.56X | 15.20X |
| **Percent Genome Covered** | 90.60% | 91.39% | 90.38% |
| **Mean Mapped Read Length** | 146.43 | 28.79 | 28.83 |
| **Average Identity** | 98.97% | 98.73% | 99.71% |
| **Mismatch Rate** | 1.30% | 1.10% | 0.82% |
| **Deletion Rate** | 0.01% | 1.25% | 1.28% |
| **Insertion Rate** | 0.00% | 0.46% | 0.48% |

Among GenoCare (K-12 reference) errors, deletion rate was highest (1.25%), followed by mismatch rate (1.10%), and insertion rate (0.46%). Among mismatch errors, C to T and G to A errors are highest (0.271% and 0.281%, respectively), each accounting for about a quarter of all mismatch errors (**Table 2**). These two most prevalent mismatch errors might be explained by a presence of thymine-guanine (C to T) and adenine-cytosine (G to A) wobbles during nucleotide incorporation in the sequencing process. Further investigation and error characterization is necessary.

**Table 2.** Distribution of mismatch error rates for GenoCare aligned to K-12 reference (total mismatch error rate: 1.10%). Gray backgrounds indicate transitions; white backgrounds indicate transversions.

|  |  |  |  |  |  |
| --- | --- | --- | --- | --- | --- |
|  |  | **To** |  |  |  |
| **From** |  | **A** | **G** | **T** | **C** |
|  | **A** |  | 0.103% | 0.081% | 0.063% |
|  | **G** | 0.271% |  | 0.036% | 0.016% |
|  | **T** | 0.026% | 0.019% |  | 0.078% |
|  | **C** | 0.103% | 0.016% | 0.280% |  |

Read length distribution for GenoCare reads mapped to the K-12 genome can be seen in **Fig. 1.** A read length peak can be observed at 20 base pairs and average mapped read length was found to be28.79 base pairs. The reported average read length can be attributed in part to the number of cycles run (120); so, there is potential for longer read length as cycle number is increased on the GenoCare platform.

**Figure 1.**
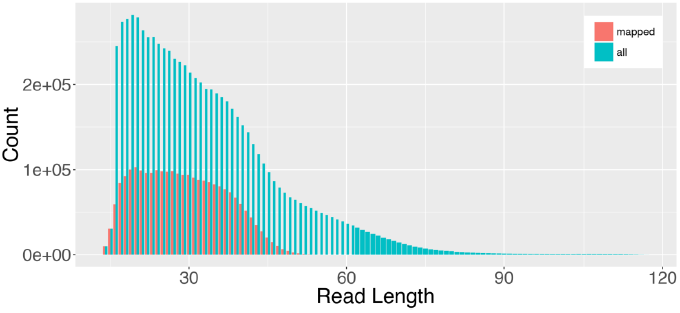
Distribution of read lengths among mapped and all filtered reads for GenoCare platform.

For GenoCare reads aligned to the K-12 genome, average coverage was 14.56X, while Illumina reads aligned to the K-12 genome had average coverage of 84.71X. Non-identity between both Illumina and GenoCare reads and the K-12 reference genome is demonstrated in low-coverage troughs (**Fig. 2)**. The highly similar depth distributions between the GenoCare and Illumina platforms reveal comparable performance. Both alignments show similar distributions of fraction of genome covered in relation to sequencing depth, with the majority of bases covered at their average depths (**Fig. 3)**.

**Figure 2.**
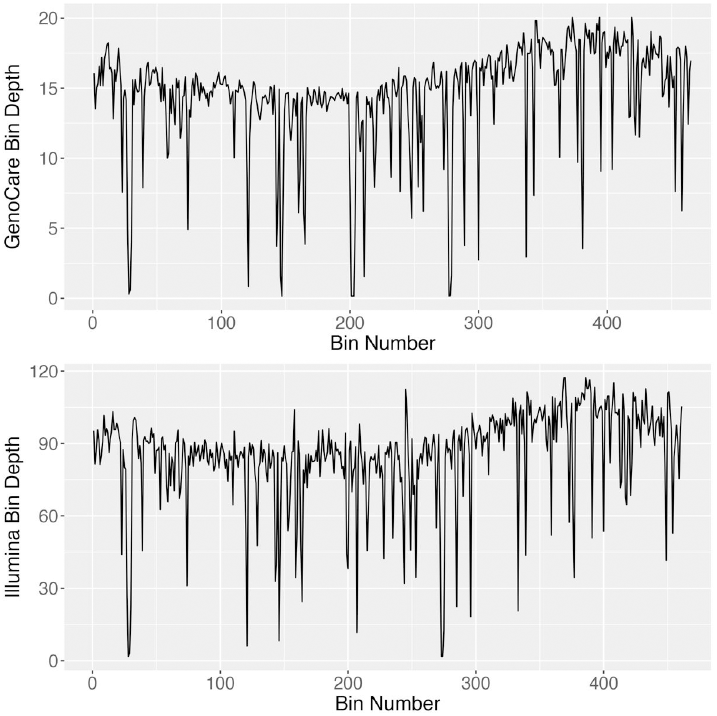
Depth distribution of GenoCare (top) and Illumina (bottom) reads by genome position, binned by 10kb.

**Figure 3.**
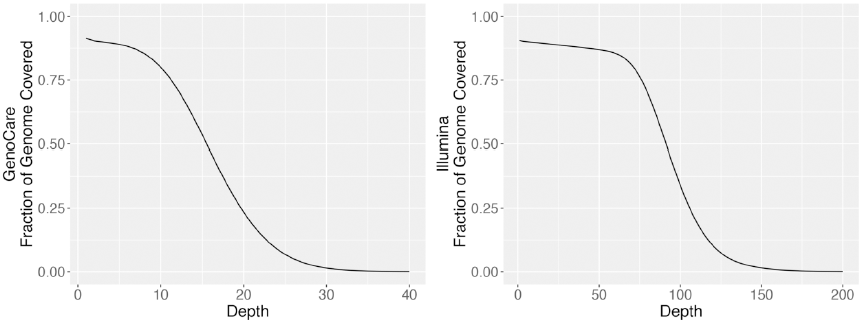
Fraction of genome covered by depth for GenoCare (left) and Illumina (right) platforms.

### CONCLUSION

In this study, we demonstrated the GenoCare platform’s capabilities for single molecule sequencing. Overall sequencing time including sample preparation and instrument run time is improved over NGS standards. This greater efficiency is critical for rapid results and diagnosis in the clinic, particularly in the advent of precision medicine. This study used only a small fraction of the GenoCare’s flow cell capacity; thus, the GenoCare is capable of far greater throughput and has potential for whole human genome sequencing. GenoCare uses poly-T oligonucleotides to hybridize with poly-A tailed DNA. This design allows for GenoCare’s potential as a new technology to handle naturally poly-A tailed RNA, and address the need for innovations in transcriptomics. GenoCare is an automated desktop sequencer for dedicated use in the clinic with potential to eclipse NGS technologies as a faster and cheaper option.

